# High-dimensional Biomarker Identification for Scalable and Interpretable Disease Prediction via Machine Learning Models

**DOI:** 10.1101/2024.10.04.616748

**Authors:** Yifan Dai, Fei Zou, Baiming Zou

**Affiliations:** Department of Biostatistics, University of North Carolina at Chapel Hill, Chapel Hill, NC, USA; Department of Genetics, University of North Carolina at Chapel Hill, Chapel Hill, NC, USA; School of Nursing, University of North Carolina at Chapel Hill, Chapel Hill, NC, USA

**Keywords:** Deep learning, Feature prescreening, Genomics data integration, Model interpretation, Nonlinear association

## Abstract

Omics data generated from high-throughput technologies and clinical features jointly impact many complex human diseases. Identifying key biomarkers and clinical risk factors is essential for understanding disease mechanisms and advancing early disease diagnosis and precision medicine. However, the high-dimensionality and intricate associations between disease outcomes and omics profiles present significant analytical challenges. To address these, we propose an ensemble data-driven biomarker identification tool, Hybrid Feature Screening (HFS), to construct a candidate feature set for downstream advanced machine learning models. The pre-screened candidate features from HFS are further refined using a computationally efficient permutation-based feature importance test, forming the comprehensive High-dimensional Feature Importance Test (HiFIT) framework. Through extensive numerical simulations and real-world applications, we demonstrate HiFIT’s superior performance in both outcome prediction and feature importance identification. An R package implementing HiFIT is available on GitHub (https://github.com/BZou-lab/HiFIT).

## Background

High-dimensional omics data, such as genomics, proteomics, and other types of biomedical data generated from high-throughput technologies, have revolutionized clinical research and personalized medicine by providing detailed molecular profiles of individuals [1–3]. Omics data offer complementary patient information in addition to low-dimensional baseline demographic and clinical features. This information helps healthcare professionals gain a deep understanding of the genetic and molecular mechanisms underlying complex human diseases, enabling improved early disease diagnoses and effective personalized treatment strategies tailored to individual patients or subpopulations [4]. However, accurately predicting disease outcomes remains highly challenging due to complex disease mechanisms, including nonlinear effects of molecular biomarkers and clinical features on disease outcomes, as well as interactive effects among these risk factors. Consequently, conventional parametric methods, such as multiple linear or logistic regression, prove ineffective in constructing powerful predictive models.

Machine learning algorithms, particularly deep neural networks (DNNs, [5]), support vector machines (SVMs, [6, 7]), random forests (RFs, [8]), and gradient boosting machines such as XGBoost [9] have demonstrated potential in handling the intricate associations between molecular biomarkers, biological, and clinical features, and disease outcomes. These approaches are adept at integrating biomedical data from various platforms to improve clinical outcome predictions, such as heart disease prediction [10] and cancer prognosis [11, 12]. However, both parametric and machine learning predictive models are complicated by the high-dimensionality of multi-omics data, commonly referred to as the “curse of dimensionality” [13–15]. Noise features in high-dimensional feature spaces often lead to poor predictive models, regardless of the modeling approach used. Specifically, predictive models with high-dimensional input features are susceptible to overfitting, which often results in high training accuracy but poor testing performance [16]. Various methods have been proposed to mitigate this issue. For example, dropout is widely utilized in DNN models to prevent the coadaptation of feature detectors [17], while pruning is another popular technique to avoid model overfitting [18]. The bootstrap bagging and scoring algorithm [19] also addresses the overfitting issue in DNN models. However, none of these adaptive methods can fully overcome the curse of dimensionality. Furthermore, as the number of input features increases, the computational complexity and workload of machine learning algorithms escalate. This escalation can lead to slow or failed convergence during the model training process, or even become computationally prohibitive. In addition, interpreting machine learning models trained on high-dimensional data presents another significant challenge. Though there are existing methods for assessing feature importance for interpretable predictions using machine learning methods, such as the Permutation Feature Importance Test (PermFIT, [20]), Shapley Additive Explanations (SHAP, [21, 22]), and Knock-off Randomized Testing [23, 24], all struggle with the curse of dimensionality. They are limited in their ability to differentiate true causal features from noise features when the data dimension is high.

To address the dimensionality issue, researchers have developed and extensively studied shrinkage linear or generalized linear models. These include methods such as the least absolute shrinkage and selection operator (Lasso, [25]), smoothly clipped absolute deviation [26], elastic net [27] and minimax concave penalty [28]. While powerful, these shrinkage algorithms suffer from performance degradation as the number of nuisance input features increases. Consequently, feature pre-screening becomes necessary. One such approach is Sure Independence Screening (SIS) [29], which selects features with strong marginal effects from high-dimensional data before applying more refined analyses like Lasso [29, 30]. Evaluating the marginal association between each input feature and a disease outcome [31, 32] can be done using metrics such as mutual information (MI), Spearman correlation, maximal information coefficient [33]), or Kendall rank correlation coefficient (Kendall’s tau). For capturing nonlinear associations, researchers may consider methods like the Hilbert-Schmidt independence criterion (HSIC [34, 35]) and kernelized partial correlation (KPC, [36]).

Despite significant advances, several challenges persist. First, the marginal screening approaches mentioned earlier are constrained by individual criteria for measuring marginal dependency, and their performance is often data-dependent [37]. A recent study [37] demonstrates that no single screening method consistently outperforms others. To address this, we propose an efficient **H**ybrid **F**eature **S**election (HFS) framework that combines multiple dependency metrics. The HFS approach identifies important features by assembling metrics, minimizing the risk of missing important features by relying only on one specific dependency measure. Additionally, we introduce a novel data-driven method that uses the isolation forest algorithm [38] to determine an optimal cutoff for dependency statistics, enabling principled identification of essential features to boost feature pre-screening performance.

While HFS significantly filters out many unimportant features, it inevitably selects some noise features due to its marginal screening nature. Therefore, a further refinement process is necessary to fine-tune the HFS list and determine the most relevant features for outcome prediction. Furthermore, it is critically important to evaluate the impact of each individual pre-selected biomarker on disease outcomes by adjusting for the confounding effects of other biomarkers under complex association settings. This deeper understanding of disease mechanisms can inform better clinical decisions. Similar to SIS, which relies on penalized linear regression for refinement, we leverage PermFIT ([20]), a computationally efficient algorithm for evaluating feature importance scores under potential complex associations using machine learning models. Combining PermFIT with HFS, we alleviate the curse of dimensionality, allowing more effective detection of complicated impacts such as nonlinear interactions among omics features on disease outcomes. We consolidate the entire process into a comprehensive high-dimensional feature importance test (HiFIT) framework. This framework encompasses feature pre-screening, refinement, and final predictive modeling, achieving robust, scalable, and interpretable outcome predictions in high-dimensional settings.

## Results

To evaluate the performance of HiFIT in eliminating nuisance features and predicting outcomes, we conducted comprehensive simulation studies under various data complexity scenarios with varying numbers of features and data generation schemes. For comparison, we included the gold standard approach, which uses only causal features as input, alongside Lasso and other machine learning algorithms. Additionally, HiFIT was applied to two real-world datasets: the weight loss data after bariatric surgery (BS) [39, 40], and RNA sequence data from three kidney cancer studies in The Cancer Genome Atlas (TCGA) [41]. Given the unknown biological ground truth of the associations between clinical and omics features and disease phenotypes, we evaluated the performance of each comparison method by focusing on the prediction accuracy of the final HiFIT model in real data applications.

## Simulation Studies

We examine the performance of the proposed methods under the following simulation scenarios: (a) **Linear case:** where the effects of the causal features are linear and additive. (b) **Non-linear case:** where the effects of the causal features are non-linear and non-additive. Specifically, we simulate *y* as follows:

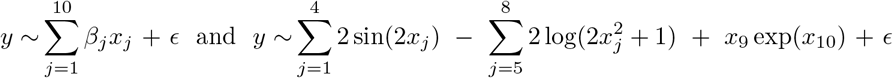

for cases (a) and (b) respectively, where **x** = (*x*_1_, …, *x*_*p*_)^*T*^ is a p-dimensional random variable drawn from a multivariate normal distribution MVN(0, *I*_*p*_), the error term *ϵ* follows a standard normal distribution, i.e., *N* (0, 1), and the vector *β* = (*β*_1_, *…, β*_10_) is drawn from a uniform distribution 𝒰 (1, 1.5) at the beginning of the simulation and remains unchanged throughout subsequent Monte Carlo simulations. For both cases, we set the total number of causal features to 10, with the remaining *p −* 10 features considered as nuisance variables. The total number of samples is set to 500. To evaluate the impact of input feature dimensions, we vary *p* across the values {500, 1000, 10000}. We split the simulated data into training and testing sets in a 9:1 ratio. For each scenario, the total number of simulation is set to 100.

We begin by comparing the performance of HFS with other feature pre-screening methods, including Lasso, Pearson Correlation (PC), Spearman Correlation (SPC), MIC and HSIC (Hilbert-Schmidt Independence Criterion). Figure 1 illustrates the ranks of correlation scores for causal features relative to all features. Since Lasso does not directly produce a correlation score, we rank the penalty parameter associated with the causal features. Specifically, we consider the largest penalty parameter at which the coefficient of a feature becomes non-zero. Under linear settings, when *p* = 500 or 1000, HFS performs comparably to parametric models like Lasso and PC in ranking causal features. As the dimensionality *p* increases to 10,000, HFS still outperforms non-parametric methods such as HSIC and MIC. Notably, HSIC is significantly influenced by noisy variables and ranks half of the causal features with smaller linear effects much lower than HFS. The advantage of HFS becomes even more pronounced in non-linear scenarios. Parametric methods like Lasso and PC become ineffective and fail to identify causal features, as shown in the second panel of Figure 1. On the other hand, while SPC and MIC relax the parametric assumptions of Lasso and PC, they only rank half of the causal features at the top. Specifically, SPC struggles to detect monotonic associations, while MIC tends to capture spurious correlations between nuisance features and outcomes. Furthermore, their performance deteriorates as the dimensionality increases to 10,000. In contrast, HFS not only identifies the largest number of causal features but also remains robust in high-dimensional data scenarios. In summary, HFS effectively combines the advantages of both parametric and non-parametric methods for feature screening by leveraging different marginal association dependencies. Other methods are either constrained by their parametric assumptions or limited in their ability to handle high-dimensional data.

**Fig. 1.**
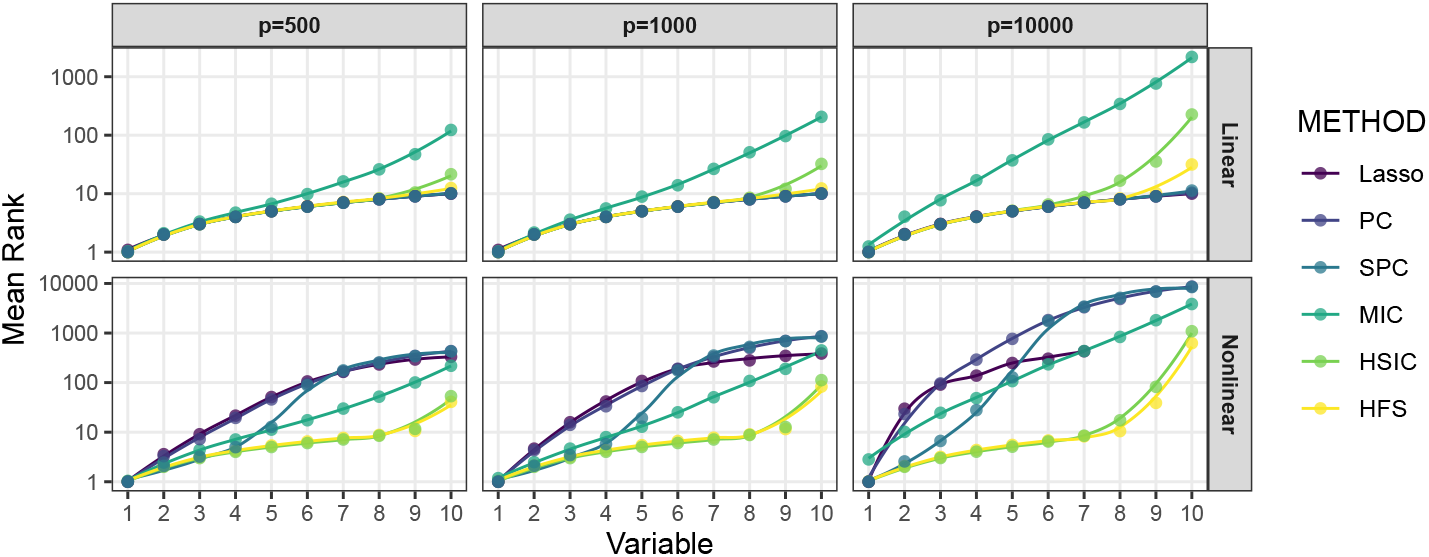
Average Rank of Causal Features Selected by Pre-Screening Methods. The x-axis denotes the number of selected causal features, and the corresponding value of the y-axis represents the average rank of this feature over 100 repetitions. The curves are generated by locally estimated scatterplot smoothing.

In practice, the cutoff for the HFS score is determined using a data-driven approach (see Methods). Features with HFS scores higher than the cutoff are retained for downstream analysis. To ensure a fair comparison between HFS and other pre-screening methods mentioned earlier, we select the same number of features across all comparison methods. We evaluate the quality of the feature list based on true positive rates (TPR) and false discovery rates (FDR). TPR is defined as the ratio of selected causal features to the total number of causal features, while FDR is defined as the ratio of selected nuisance features to the total number of selected features. Figure 2a illustrates the quality of feature sets. HSIC and MIC tend to overlook causal features with smaller effects under linear settings. Meanwhile, Lasso, PC, and SPC struggle to detect log-quadratic or interaction terms. In comparison, HFS consistently identifies more causal features across all simulation scenarios. However, as the dimensionality increases, the FDR of all pre-screening methods inflates. To address this issue, HiFIT refines the pre-screening feature set obtained from HFS using machine learning algorithms. Figure 2b compares (i) HFS feature sets with the cutoff parameter determined by XGB, RF, SVM, and DNN (denoted as S-XGB, S-RF, S-SVM, and S-DNN); (ii) HiFIT feature sets obtained by applying PermFIT to the corresponding HFS feature lists, retaining features with p-values smaller than 0.1 (denoted as HF-XGB, HF-RF, HF-SVM, and HF-DNN).

**Fig. 2.**
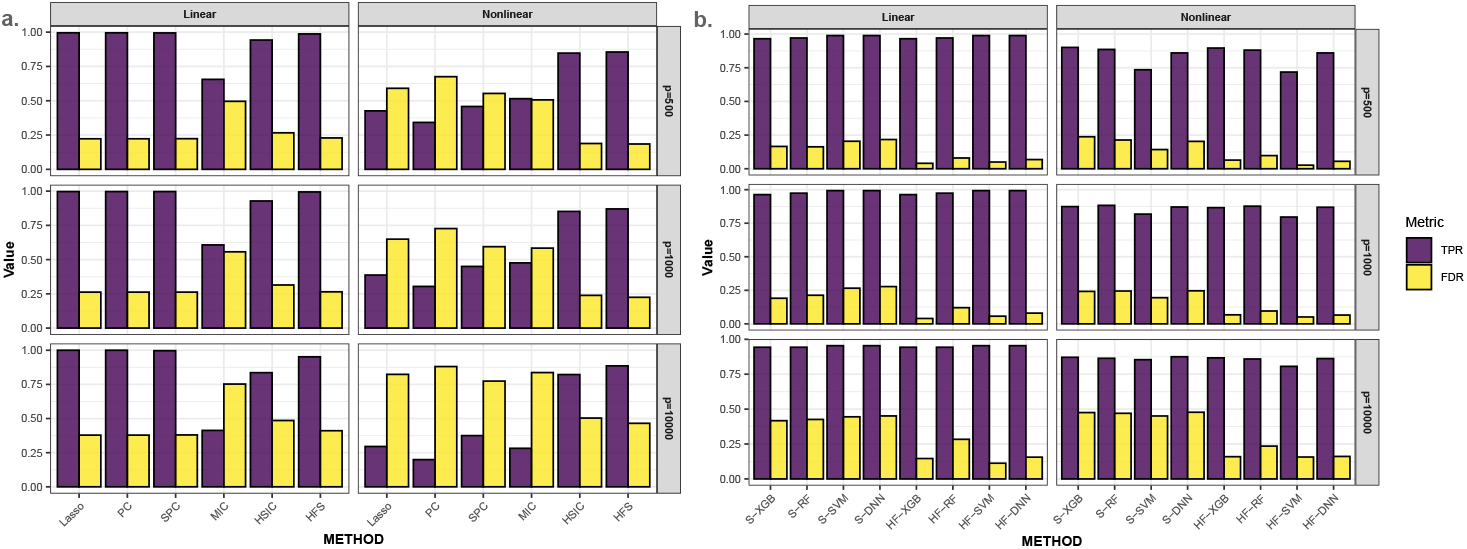
High-dimensional Feature Pre-screening and Selection Results. **(a)** Performance of feature pre-screening methods. **(b)** Feature selection results of HiFIT models. TPR and FDR are averaged over 100 simulations.

HiFIT feature sets retain most of the causal features identified by HFS. Specifically, the TPRs of HF-XGB, HF-RF, and HF-DNN match those of the corresponding HFS feature sets, and the TPR of HF-SVM is comparable to S-SVM. Furthermore, HiFIT improves the FDR of HFS. For instance, when *p* = 500 or 1000, the FDR of HF-XGB, HF-SVM, and HF-DNN is controlled at 0.1. As *p* increases to 10,000, HF models reduce the FDR of the corresponding HFS sets from 50% to less than 20%, and further reductions are possible with p-value adjustments.

Next, we compare the importance scores and p-values from HiFIT with those from PermFIT. Figure 3 provides detailed feature importance scores and corresponding p-values for the following scenarios: (i) HiFIT models: HF-DNN, HF-SVM, HF-XGB, and HF-RF; and (ii) PermFIT models: PermFIT-DNN, PermFIT-SVM, PermFIT-SVM, and PermFIT-XGB (applied to the same models without pre-screening). We observe that all HiFIT models successfully identify causal features, consistently estimating high importance scores across various simulation scenarios. HiFIT assigns low scores and large p-values to nuisance features, demonstrating its ability to control type-I errors reasonably in high-dimensional settings, regardless of association complexity. In contrast, PermFIT models struggle with nonlinear effects. PermFIT-DNN and PermFIT-SVM identify only one nonlinear causal feature out of ten on average. While PermFIT-RF and PermFIT-XGB successfully identify all nonlinear features, their importance scores for causal features are substantially lower than those from HF-XGB and HF-RF, highlighting HiFIT’s superior efficiency. For instance, the average p-value of the interaction term *X*_10_ from PermFIT-RF is higher than that from HF-RF, indicating that PermFIT-RF is more likely to overlook this feature. As data dimensionality increases, PermFIT’s computation cost becomes overwhelming when *p* = 10, 000, whereas HiFIT remains scalable to high-dimensional data.

**Fig. 3.**
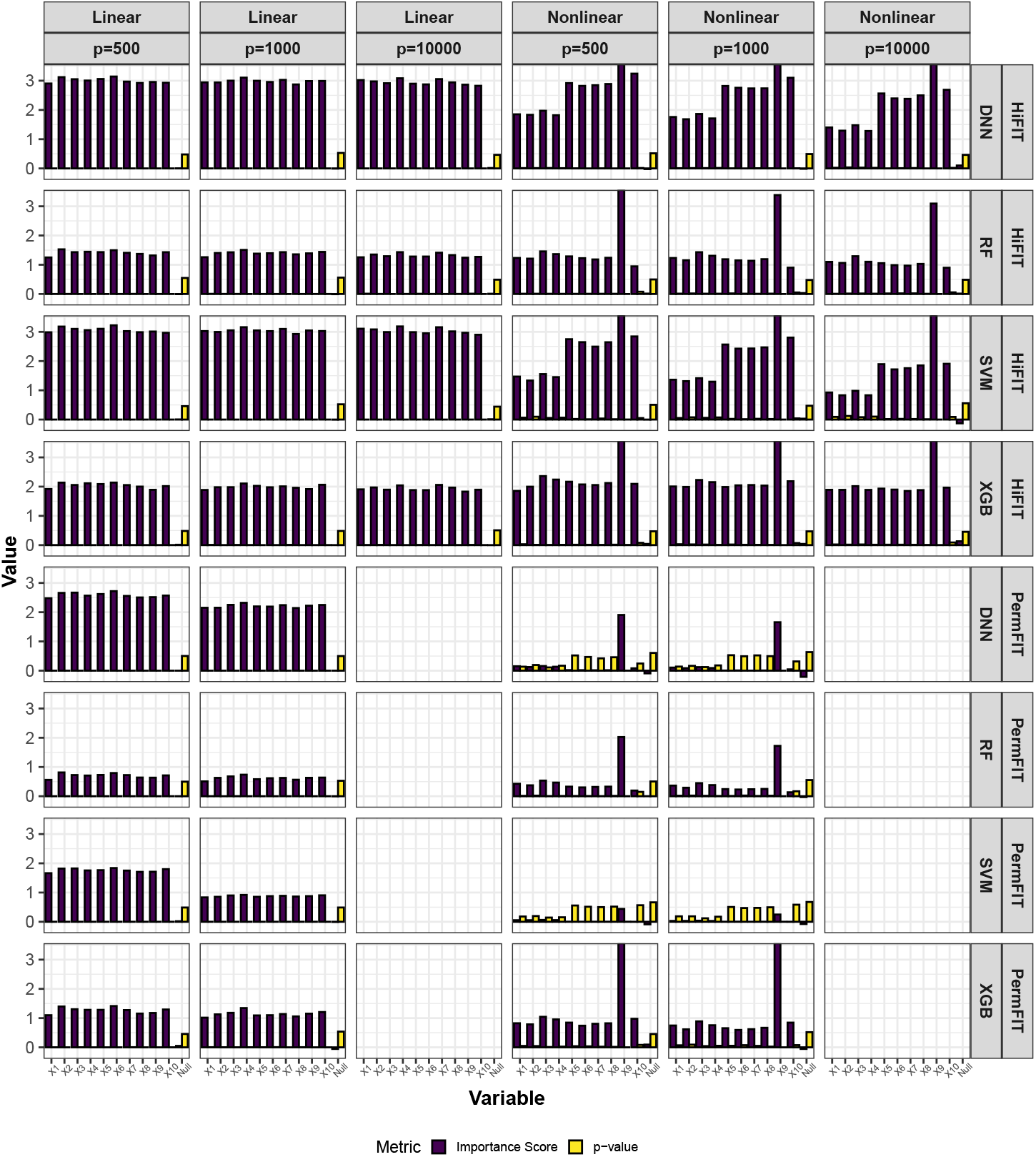
HiFIT Feature Interpretation Results. Average feature importance scores and p-values for 10 causal features (denoted as *X*_1_, …, *X*_10_) and the feature set of nuisance features (denoted as null) over 100 repetitions. Importance scores of features not selected by HFS are set to zero.

Finally, we assess the impact of HFS and HiFIT on prediction accuracy for various machine learning models. Figure 4 summarizes the predicted average mean squared error (MSE) and Pearson correlation coefficient (PCC) for the following scenarios: (i) Taking the full features as input: Denote the corresponding models as Lasso, DNN, XGB, RF, and SVM; (ii) Taking the HFS pre-screened features as input: Denote the corresponding models as S-Lasso, S-DNN, S-XGB, S-RF, and S-SVM; (iii) Taking the HiFIT refined features (with *p ≤* 0.1) as input: Denote the corresponding models as HF-DNN, HF-XGB, HF-RF, and HF-SVM. Under linear settings, and for scenario i), all machine learning models perform worse than Lasso, and their prediction errors increase with dimensionality. When *p* = 10, 000, RF, DNN, and SVM all fail to converge. After HFS, S-SVM and S-DNN achieve performance comparable to Lasso across dimensions, with similar PCC and MSE. S-RF and S-XGBoost perform slightly worse than Lasso but close to their optimal predictions (using only true causal features). Across dimensions, models trained with HFS features exhibit PCC no less than 5% below their counterpart gold models (except for Lasso in nonlinear settings due to model misspecification). Without pre-screening, all four machine learning methods struggle to make reliable predictions (see Figure 4). SVM and DNN even have higher MSE than Lasso. Trained with the only 10 causal features, SVM and DNN outperform XGBoost and RF, highlighting their better ability in capturing complex feature-outcome relationships (see Table 1) when a set of right features are provided. However, in high-dimensional settings with many nuisance features, their performance lags behind Lasso, RF, and XGBoost, and they even become computationally infeasible for *p* = 10, 000. In contrast, HFS significantly improves performance of these models, especially for SVM and DNN. S-DNN achieves the lowest prediction error and highest PCC across dimensions, surpassing other machine learning algorithms. Even at *p* = 10, 000, S-DNN maintains a high PCC of 0.77. Overall, HFS enhances machine learning models by reducing MSE (by 30%) and increasing PCC (by 20%) for *p* = 500 and 1000. The models remain stable as feature dimension increases, with PCC consistently above 0.7.

**Table 1.** Prediction Performance of HiFIT Models.

| Setting | Model | p=10* |  | p=500 |  | p=1000 |  | p=10000 |  |
| --- | --- | --- | --- | --- | --- | --- | --- | --- | --- |
|  |  | MSE | PCC | MSE | PCC | MSE | PCC | MSE | PCC |
| Linear | XGBoost | 4.70 | 0.85 | 4.84 | 0.84 | 4.87 | 0.85 | 5.51 | 0.82 |
|  | RF | 5.54 | 0.90 | 5.89 | 0.89 | 6.21 | 0.89 | 7.82 | 0.85 |
|  | SVM | 1.03 | 0.97 | 1.21 | 0.96 | 1.14 | 0.96 | 1.28 | 0.96 |
|  | DNN | 1.12 | 0.96 | 1.30 | 0.96 | 1.27 | 0.96 | 1.43 | 0.95 |
| Nonlinear | XGBoost | 9.41 | 0.81 | 10.57 | 0.77 | 10.78 | 0.77 | 12.33 | 0.72 |
|  | RF | 12.94 | 0.79 | 13.87 | 0.76 | 14.58 | 0.74 | 15.03 | 0.70 |
|  | SVM | 9.25 | 0.80 | 11.42 | 0.74 | 12.94 | 0.69 | 13.98 | 0.66 |
|  | DNN | 5.01 | 0.91 | 6.78 | 0.87 | 7.41 | 0.85 | 10.38 | 0.77 |

\* Oracle model including 10 causal features only as input features

**Fig. 4.**
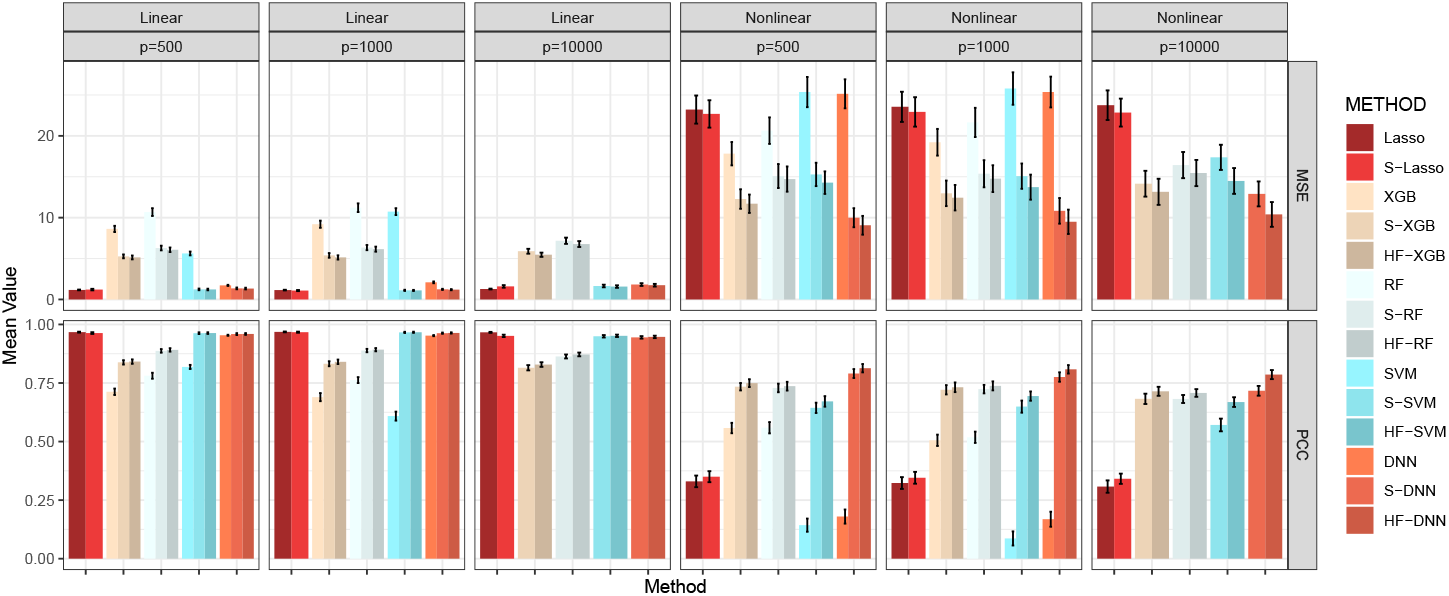
Average MSE and PCC for Methods in Comparison. Lasso, XGB, RF, SVM, and DNN: specific models with all features; S-Lasso, S-XGB, S-RF, S-SVM, S-DNN: specific models with HFS pre-screening; HF-XGB, HF-RF, HF-SVM, HF-DNN: specific models with HiFIT feature selection. Simulation in each scenario is repeated 100 times.

In addition to the comprehensive simulation studies conducted for continuous outcomes, we also conducted extensive numerical studies to evaluate the performance of HiFIT for binary data. Similar conclusions were observed, and detailed results are presented in the supplementary materials. Beyond the simulation studies, we applied the proposed HiFIT framework to two real-world applications.

### Weight Loss After Bariatric Surgery Study

In the first real application, we applied the HiFIT to a weight loss study for bariatric surgery (BS). One of the primary objective of this study was to use the baseline microbiome profiles along with other demographic and clinical features to predict postoperative weight loss and identify associated important features. The weight loss microbiome cohort consists of 144 participants undergoing bariatric surgery with 50% of them having Roux-en-Y Gastric Bypass (RYGB) and the other 50% having Sleeve Gastrectomy (SG) [39, 40]. The body mass index (BMI) and fecal material were collected from individuals at 1, 6, 12, 18, and 24 months post-surgery. The microbial profiles of the BS study were characterized through shotgun Whole Genome Sequencing across multiple time points of 135 subjects (*n* = 135), resulting in total 430 measurements of BMI change with 1533 microbial genera. There are also four demographic features of participants, e.g., age, race, height, and sex. In this study, we aim to predict patients’ BMI change from the day of surgery to their last recorded measurement, using their surgery type, time since surgery, demographic features, and the gut microbiome abundance collected from their first visit. Since the effect of the surgery type and demographic features on weight change is of particular interest [39], we retain all these features and only perform pre-screening on the microbiome features with HFS. Figure 5 presents the model performance, demonstrating that HFS substantially enhances the prediction accuracy of Lasso, SVM, RF, and DNN. HiFIT further improves the performance of RF and DNN. Specifically, HF-DNN achieves the highest prediction accuracy among all methods, with the smallest predicted MSE and the largest PCC.

**Fig. 5.**
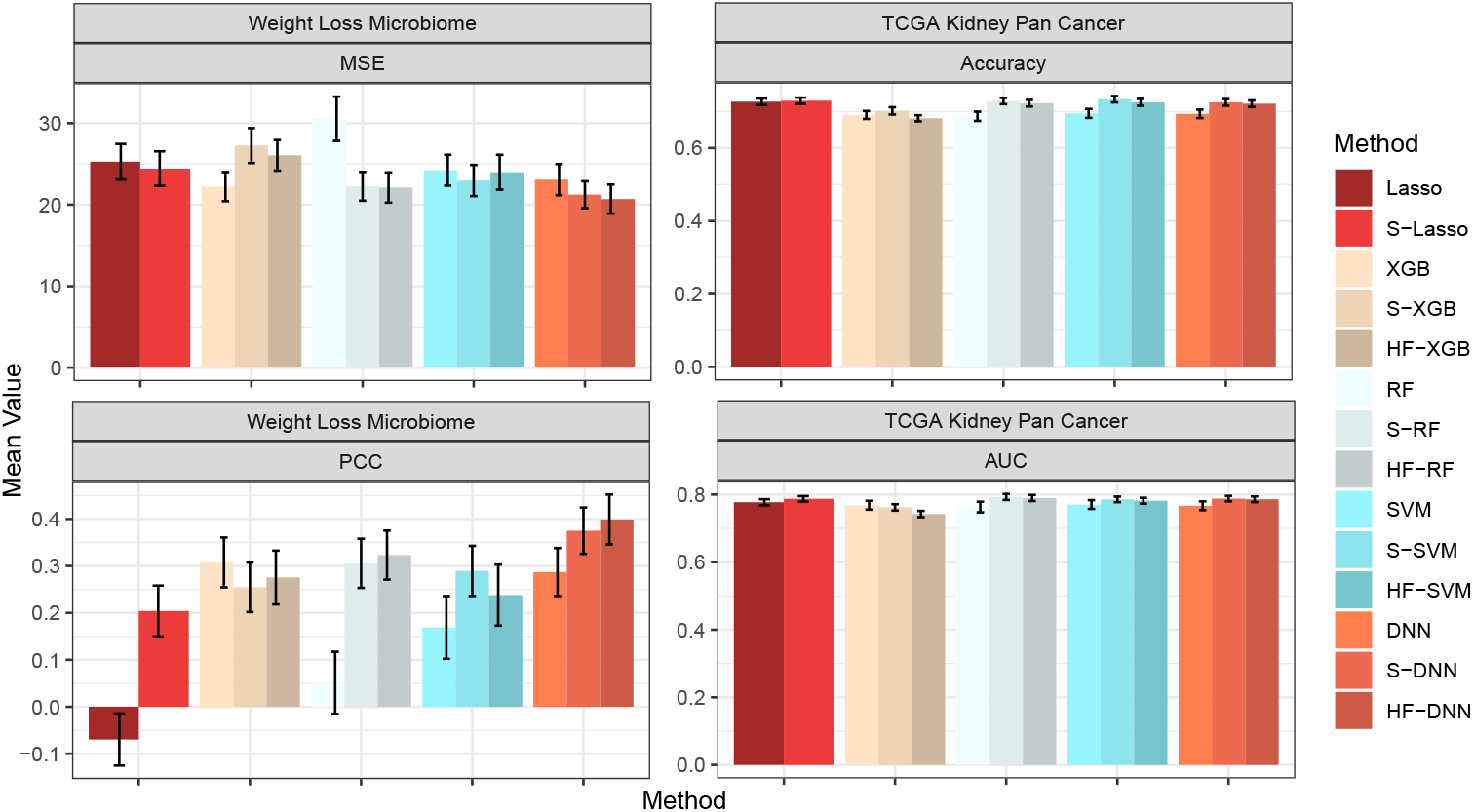
Model Performance for the Real Data Analysis. The first column presents the MSE and PCC for the weight loss microbiome study. The second colume presents the accuracy and AUC for TCGA kidney pan cancer cohort. All metrics are averaged on separate testing sets consisting of 10% observations with random repeats for 100 times.

Figure 6a presents the feature-wise p-values from the four machine learning models from HiFIT. Some demographic features, including time, age, and race, have significant effects on weight loss after BS. This finding on age is consistent with a previous study by Contreras et. al. [42]. HiFIT provides further insights into microbial effects on weight loss. As shown in Figure 6a, all four machine learning models identify important microbiota on weight loss. This implies that gut microbial abundance offers a distinct source of information on weight loss beyond patients’ demographic features. Specifically, all HiFIT models identify *Hyphobacterium* as an important predictor for post-BS weight loss, and three HiFIT models highlight *Panacibacter* as an important genus.

**Fig. 6.**
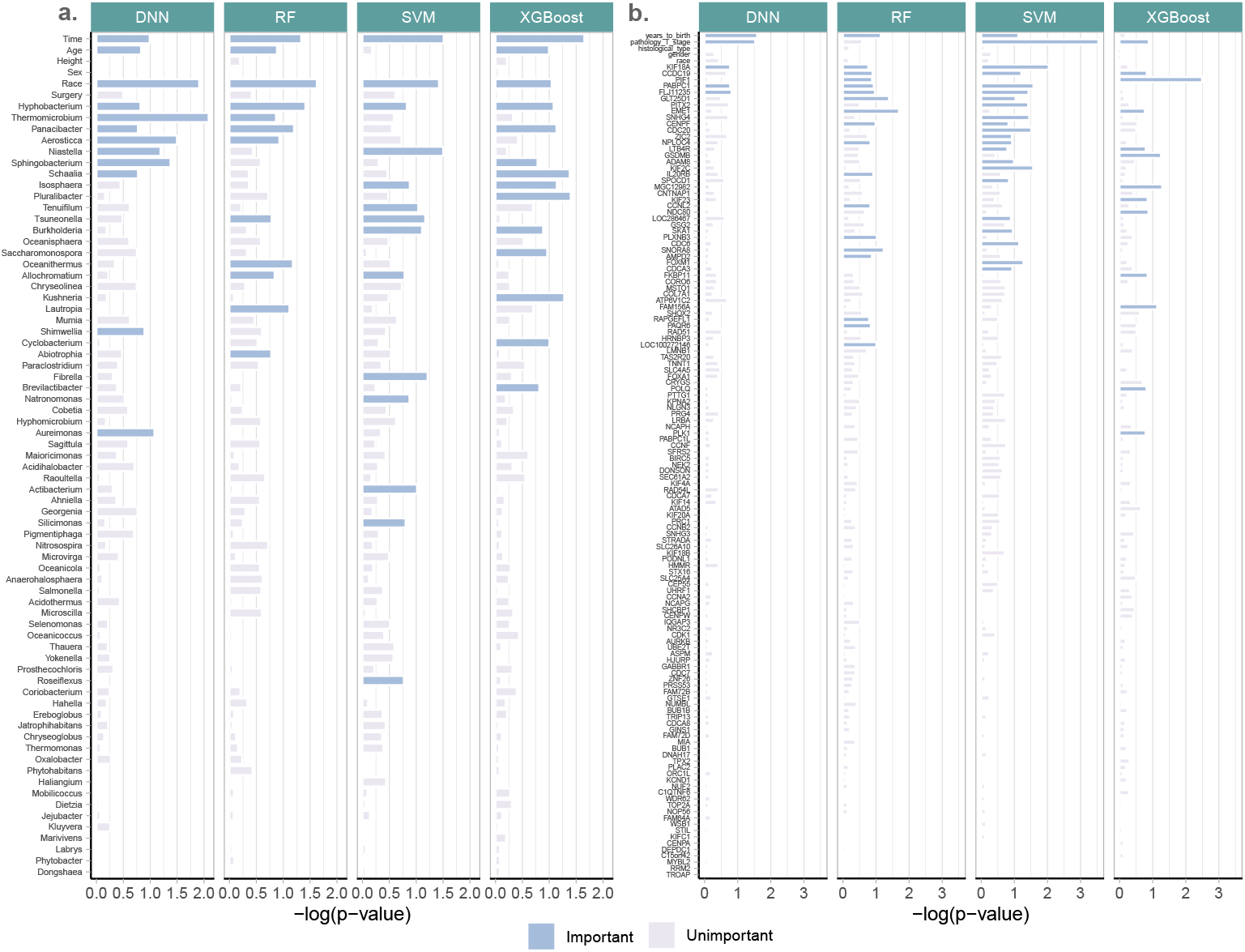
Negative log_20_ p-values for biomarkers from real datasets. **(a)** Feature importance for the weight loss data. **(b)** Feature importance for the TCGA data.

### TCGA Kidney Cancer Data from the KIPAN Cohort

We further applied the HiFIT to analyze binary outcomes using TCGA data. Though TCGA has a large collection of publicly available clinical and omics data [41], we focus on the Pan-kidgeny corhort (KIPAN, *n* = 941) in our analysis. We aim to predict the patients’ survival status using normalized counts of RNA sequence data from Illumina HiSeq platform at gene level. For simplicity, we categorize patients into two groups: long-term survival (≥ 5 years of survival) and short-term survival (*<* 5 years of survival). Out of the total 941 samples in this cohort, 193 participants achieved long-term survival, while 242 participants achieved short-term survival. The remaining samples were lost to follow-up. Additionally, expression profiles from 20,189 genes and five clinical features — age, tumor stage, histological type, gender, and race — are available for these patients. Due to the large number of features and the limited sample size in the cohort, RF, SVM, and DNN fail to converge when using the full set of features, making reliable predictions impossible. Instead, we implement these algorithms using the top 2000 genes as training input. Figure 5 presents the comparison of prediction accuracy and AUC using HFS/HiFIT selected genes. HFS improves the performance of all four machine learning algorithms and Lasso. Specifically, S-RF yields the highest prediction accuracy and AUC among all methods. Although HiFIT does not further improve the performance of machine learning algorithms, the prediction accuracy and AUC of HF-RF, HF-SVM and HF-DNN are still comparable to S-RF, S-SVM, and S-DNN, outperforming models using all top 2000 differentially expressed genes as input. Despite the minor improvement in prediction accuracy, HiFIT offers feature importance evaluation with solid statistical inference, offering in-depth understanding of disease mechanisms.

The feature-wise p-value for kidney cancer-associated features is summarized in Figure 6b. We next summarize important features that are identified by at least three models from HF-DNN, HF-RF, HF-SVM, and HF-XGB. First, age and tumor stage are two important demographic features. Second, we uncover the following important genes: *KIF18A, CCDC19*, and *PABPC1*, which are consistent with previous cancer studies. Specifically, *KIF18A* is required for chromosomally unstable tumor cells for proliferation [43] and exhibits association with pan cancer survival across multiple cohorts [44]. *PABPC1* was shown to promote cell proliferation and metastasis in pancreatic cancer [45]. A previous study [46] also identified *CCDC19* as a candidate tumor suppressor. Lastly, HF-DNN, HF-RF and HF-SVM detect one novel gene – *FLJ11235* – whose role in pan cancer deserves further investigation.

## Discussions

High-dimensional omics profiles, coupled with low-dimensional biological and clinical features, significantly influence the onset and severity of complex human diseases. To gain fundamental insights into disease mechanisms and enhance early diagnosis and precision medicine, identifying critical molecular biomarkers and clinical risk factors is essential. However, the curse of high-dimensionality, combined with intricate associations between disease outcomes and omics profiles, poses substantial analytical challenges.

In this study, we address the high-dimensionality curse by first introducing an ensemble and data-driven biomarker identification tool, i.e., HFS, which constructs a candidate feature set for downstream predictive models. We then leverages pre-screened features from HFS and further refines them using HiFIT, a computationally efficient permutation-based feature importance test. Our research demonstrates HiFIT’s superior performance through comprehensive numerical simulations and two real data applications. By effectively handling high-dimensional data and capturing complex associations among molecular biomarkers, biological and clinical features, and disease outcomes, HiFIT contributes to robust feature importance identification. This, in turn, leads to more accurate outcome predictions under high-dimensional settings.

In summary, the proposed HiFIT bridges the gap between high-dimensional omics profiles and low-dimensional biological and clinical features. It enables robust, scalable, and interpretable disease outcome predictions, providing a valuable framework for improved disease management and personalized treatment strategies.

## Methods

The proposed HiFIT framework comprises two critical components: feature prescreening by HFS and machine learning-based feature importance testing using the PermFIT algorithm [20]. HFS pre-screens high-dimensional features by evaluating their complex marginal association with the outcome, addressing the curse of dimensionality commonly encountered in biological data. After feature pre-screening, the features selected by HFS are fed into machine learning models—such as DNN, RF, XGBoost, and SVM—to build initial predictive models. PermFIT then assesses the impact of each HFS-selected feature on the outcome under intricate associations. This helps to further filter out unimportant features and build the final predictive model.

### HFS: Hybrid Feature Screening

Let **y** = (*y*_1_, *…, y*_*n*_)^*T*^ be the clinical outcome of interest across *n* samples, and **X** = {*x*_*ij*_} be an *n×p* matrix, where *x*_*ij*_ is the *j*^*th*^ feature of sample *i*, and *p* is the number of input features. Further, define **x**_*i·*_ = (*x*_*i*1_, *…, x*_*ip*_)^*T*^ and **x**_*·j*_ = (*x*_1*j*_, *…, x*_*nj*_)^*T*^, and assume that among the *p* input features, there exists a causal feature subset *S* with |*S*| *< p* such that 𝔼 [*y*_*i*_|**x**_*i·*_] = 𝔼 [*y*_*i*_|{*x*_*ij*_, *j* ∈ *S*}]. To ensure the selection power of causal features, we follow the sparse assumption of SIS such that the true number of causal features |*S*| ≪ *p*. We also follow SIS to refer the causal features as important features while the other features are referred to as noise features. In contrast to SIS, we aim to screen important features whose effects on the outcome are linear or more complex than linear by employing multiple marginal utility metrics. Specifically, HFS combines the adjusted R-square from polynomial regression with kernel partial correlation (KPC, [36]) coefficient. It utilizes isolation forest [38] to assign an anomaly score to each feature, where a higher anomaly score indicates a stronger association of the feature with the outcome. For the *j*^*th*^ feature, we fit the following linear or generalized linear model as

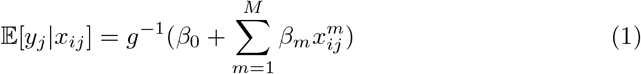

with an appropriate link function *g* corresponding to the outcome type, e.g., *g*(*x*) = *x* for continuous outcome, and *g*(*x*) = logit(*x*) for binary outcome, and a pre-specified order *M* polynomial. The adjusted R-square or McFadden’s pseudo R-square [47], denoted as *ρ*_*j*1_, is computed as follows:

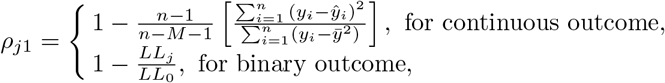

where 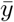 is the sample mean of the outcome, and ŷ _*i*_ is the predicted outcome of sample *i*; *LL*_*j*_ is the maximal log-likelihood of model (1), and *LL*_0_ is the maximal log-likelihood of the null model with an intercept only.

For more general marginal non-linear association detection, we also compute the KPC coefficient of each feature within the Reproducing Kernel Hilbert Space (RKHS) framework ([36]) where the *n × n* sample kernel matrices are defined as:

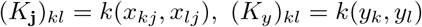

The centered kernel matrices are denoted as 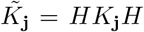 and 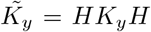 where 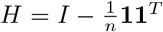. Here, *I* denotes an *n* by *n* identity matrix, and **1** denotes a 1-vector of size *n*. The empirical KPC between *y* and the *j*^*th*^ feature is:

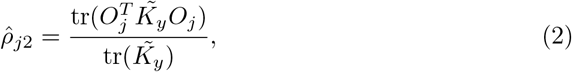

where 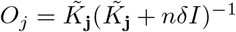 with *δ* being a positive constant. In our analysis, we use the Gaussian kernel 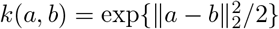.

We combine the two utility functions of the *j*^*th*^ feature into a correlation vector ***ρ***_*j*_ = (*ρ*_*j*1_, *ρ*_*j*2_)^*T*^. We then model the distribution of ***ρ***_*j*_s through a mixture model by assuming that the vector corresponding to the noise features forms one cluster with a distribution of *g*_0_ while the vector associated with the remaining ones forms another cluster with a distribution of *g*_1_. More specifically, we have

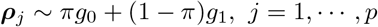

where *π* is the proportion of the noise features. The cumulative distribution function of *g*_0_, 𝔾_0_(*x*), is assumed to be no smaller than 𝔾_1_(*x*), the cumulative distribution function of *g*_1_. The features classified to the second cluster are considered anomaly features which are expected to overlap largely with set *S* as long as the marginal utilities of the important features are not too small. We utilize the isolation forest [38], a bootstrapping ensemble of isolation trees to estimate the anomaly probability of each feature as follows: for *B* bootstrap replicates, the anomaly score of feature *j* is given by

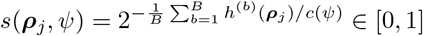

where *Ψ* is the bootstrapped sample size regularized by *c*(*Ψ*), and *h*^(*b*)^(*·*) represents the path length from the root to the leaf node that the sample belongs to, adjusted by the average path length of the *b*-th isolation tree. For a given cutoff *τ*, the selected feature set is given by

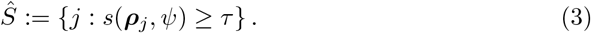

### Machine Learning Based Feature Importance Test

In this section, we outline the PermFIT procedure for assessing the importance of each individual feature. PermFIT [20] is a permutation-based feature importance score test that was originally developed for seemingly black-box machine learning models. Suppose *y* = *f* (**x**)+*ϵ*, and 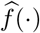 is the estimated function of *f* (·) from a machine learning model. We use some notation shortcuts and assume that the input feature vector now contains only the features in the HFS list *Ŝ*. Consequently, *p* now refers to the length of *Ŝ*, which is much smaller than the original number of high-dimensional input features. In this paper, we focus on four popular machine learning algorithms, although the framework is not limited to them: (1) SVM - implemented via R package “e1071” using Radial kernels, (2) RF - implemented via R package “randomForest”, (3) XGBoost - implemented via R package “xgboost”, and (4) ensemble DNN - implemented via R package “deepTL”. Notably, the ensemble DNN differs from conventional DNN models. It leverages a bagging and filtering algorithm to identify and tease out poorly performing bootstrapped DNN models, thereby enhancing the stability of the final DNN ensemble model [19].

Following PermFIT, for the *j*^*th*^ feature in *Ŝ*, we define its impor-tance score Λ_*j*_ as 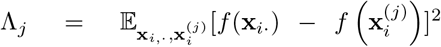, where 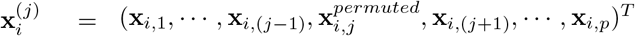 with 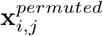 being the *i*^*th*^ element of the permuted vector of **x**_*·j*_. The importance score Λ_*j*_ equals zero when the contribution of the *j*^*th*^ feature to *y* is null, and that, the stronger the impact of the *j*^*th*^ feature on the outcome, the larger Λ_*j*_ is. PermFIT then empirically estimates Λ_*j*_ by 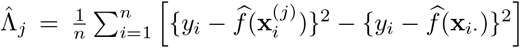. To avoid potential overfitting of *f* for data with finite samples, PermFIT employs a data splitting strategy in which it divides the data into training and validation sets. It uses the training set for generating 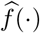 and the validation set for evaluating the distribution of Λ_*j*_. That is, let 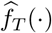 denote the estimate of *f* (·) from a training set, and 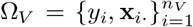 be the validation set, we obtain the feature importance score estimate 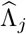, and its associ-ated variance as 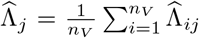 and 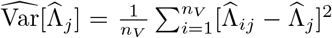 with 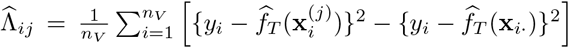. The statistics for feature importance test of *X*_*j*_ can be constructed as

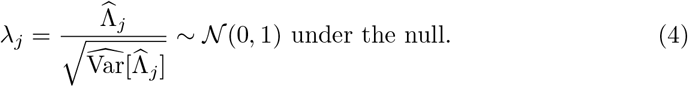

With the proposed test (4), *Ŝ* is further refined to a final feature list, denoted as *Ŝ*_*f inal*_ and used by HiFIT to build the final predictive model. The importance score test for the binary outcome can be constructed in a similar manner, and the details can be found in [20].

### Adaptive Selection of the HFS Cutoff Parameter

In this section, we propose a data-driven approach that heuristically and computationally efficiently searches for an optimal *τ* from a set of candidates (ordered from the smallest to the largest) (*τ*_0_, *τ*_1_, …, *τ*_*R*_), with {*Ŝ*_1_, …, *Ŝ*_*R*_} being the corresponding selected feature sets. These sets consist of features with their anomaly scores falling in the intervals of the candidate cutoffs. That is,

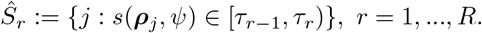

For the *r*^*th*^ feature sets, we estimate its set importance score 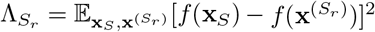, analogous to the way that PermFIT defines feature importance score, where **x**_*S*_ denotes the features with the HFS anomaly score larger than *τ*_0_, and 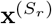 is a rearranged **x**_*S*_ with features in *S*_*r*_ replaced by random permutations. For computational efficiency, instead of testing the importance of each feature, we estimate the set importance scores here. The smallest *τ* among all candidates, for which the feature sets have a p-value smaller than 0.1 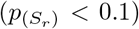, will be selected by HiFIT as the final cutoff. In this paper, we set the candidate list as (0.5, 0.55, 0.6, 0.65, 0.7) for both simulation studies and real data analysis.

### Bias Control and Cross-validation of HiFIT

As mentioned above, to avoid potential biases from the selection of *τ* and model over-fitting, HiFIT employs a nested *K*-fold cross-validations. First, it splits the training samples into *K* folds, denoted as {*𝒟*_1_, …, *𝒟*_*K*_}. One of them, say 𝒟_1_, will be selected for validation and tuning, while predictive models and HFS scores will be built and estimated with the remaining sets. The validation fold 𝒟_1_ will be further split into two subsets, 𝒟_1,1_ and 𝒟_1,2_, where 𝒟_1,1_ will be first used to tune *τ*, while the final feature importance score will be estimated with 𝒟_1,2_. Then we repeat the last two steps by switching the roles of 𝒟_1,1_ and 𝒟_1,2_. After repeating the above procedure *K* times, we estimate the mean and variance of the null distribution of the importance score of each feature and based on which we formally test the importance of each feature as done in Mi et. al [20]. Of note, different features from each cross data may be selected, and HiFIT merges all selected features into a final feature set.

### Setup for Simulation Studies and Real Data Analysis

In the simulation study, to compare the performance of HFS and other screening methods (Lasso, MIC, PC, SPC, and HSIC), we randomly split the data into a training set and a validation set at a ratio of 9:1. HFS is first implemented on the training set, and then the cutoff is determined on the validation set. The other methods are implemented on the full dataset, with the number of screened features being the same as HFS. In the simulation study and real data analysis sections, to compare HiFIT predictive models with corresponding baselines, we randomly split the data into a training set and a testing set at a ratio of 9:1. SVM, RF, XGB, and DNN are trained on the full training set, while S-SVM, S-RF, S-XGB, and S-DNN further use 1/9 of the training set to fine-tune the HFS cutoff. HF-SVM, HF-RF, HF-XGB, and HF-DNN conduct the 5-fold cross-validation on the training set as described above, with 100 permutations and the p-value cutoff set to 0.1. Lasso is implemented using the R package “glmnet” with 10-fold cross-validation to select its hyperparameter, while S-Lasso performed HFS with cutoff fixed at 0.6. The performance of the above predictive models is evaluated on the testing set. The above procedures are repeated for 100 times. The feature importance scores and p-values of two real datasets are computed by HiFIT on the full dataset with 5-fold cross-validation.

## Supplementary information

## Acknowledgement

This study was partially supported by National Institutes of Health R56 (1R56LM013784-01A1) and R01 (R01LM014407) grants.

## Authors’ contributions

Y.D. implemented the algorithms “HiFIT” for the proposed method and performed numerical analyses. All authors contributed to the methodology development and writing the manuscript.

## Availability of data and materials

The metagenomic sequences used in this study can be found at the National Center for Biotechnology information Sequence Read Archive (https://www.ncbi.nlm.nih.gov/sra) under BiopProject PRJNA668357 and PRJNA668472. The TCGA dataset is publicly available at the LinkedOmics website (http://linkedomics.org), where the KIPAN study (i.e., KIRC, KICH, and KIRP studies) is used in our analysis.

## Code availability

HiFIT is implemented in our R package “HiFIT” which is available along with source code for replicating the simulation studies and real data applications on GitHub (https://github.com/BZou-lab/HiFIT).

## Declarations

### Ethics approval and consent to participate

Not applicable.

### Consent for publication

Not applicable.

