## Supplement for "High-dimensional Biomarker Identification for Scalable and Interpretable Disease Prediction via Machine Learning Models"

### Simulation Studies on Binary Outcomes

We conducted simulation studies on binary data generated from the following models, i.e., one for generalized linear impacts and the other for generalized nonlinear effects.

$$
\begin{aligned}
\text{logit}[P(y|\mathbf{x})] &= \sum_{j=1}^{10} \beta_j x_j \\
\text{logit}[P(y|\mathbf{x})] &= \sum_{j=1}^{4} 4\sin(2x_j) - \sum_{j=5}^{8} 4\log(2x_j^2+1) + 2x_9\exp(x_{10}) + 13
\end{aligned}
$$

where $\beta_j \sim \mathcal{U}(2,3)$. All other data structures are generated in the same way as in the continuous scenarios. Results are presented in Supplementary Figures 1 $\sim$ 4. Similar to the results for continuous outcomes, HFS effectively selects most causal features, while HiFIT further reduces noise features, thereby enhancing the prediction accuracy of machine learning models.

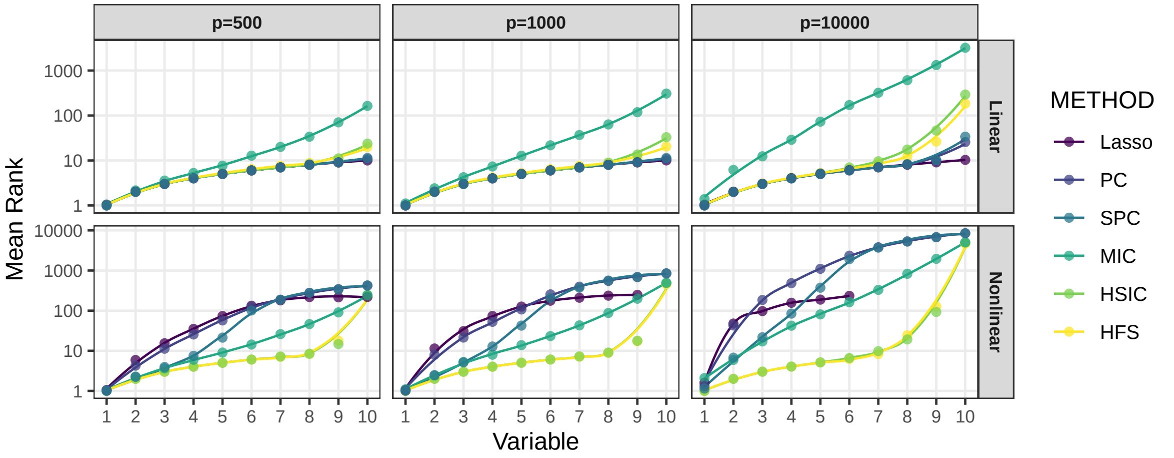


Supplementary Figure 1: **Average Rank of Causal Features Selected by Pre-Screening Methods on Binary Outcomes.**The x-axis denotes the number of selected causal features, and the corresponding value of the y-axis represents the average rank of this feature over 100 repetitions. The curves are generated by locally estimated scatterplot smoothing.

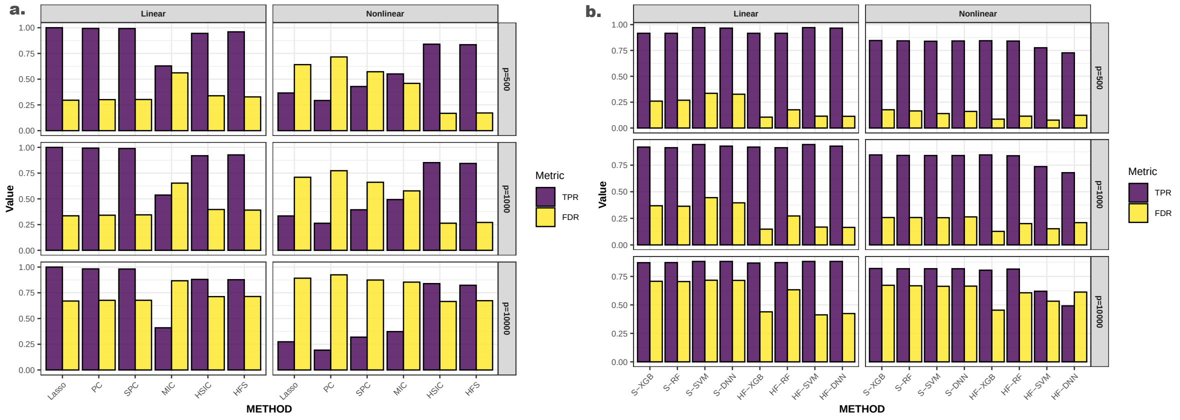


Supplementary Figure 2: **High-dimensional Feature Pre-Screening and Selection Results on Binary Outcomes.** **(a)** Performance of feature pre-screening methods. **(b)** Feature selection results of HiFIT models. TPR and FDR are averaged over 100 simulations.

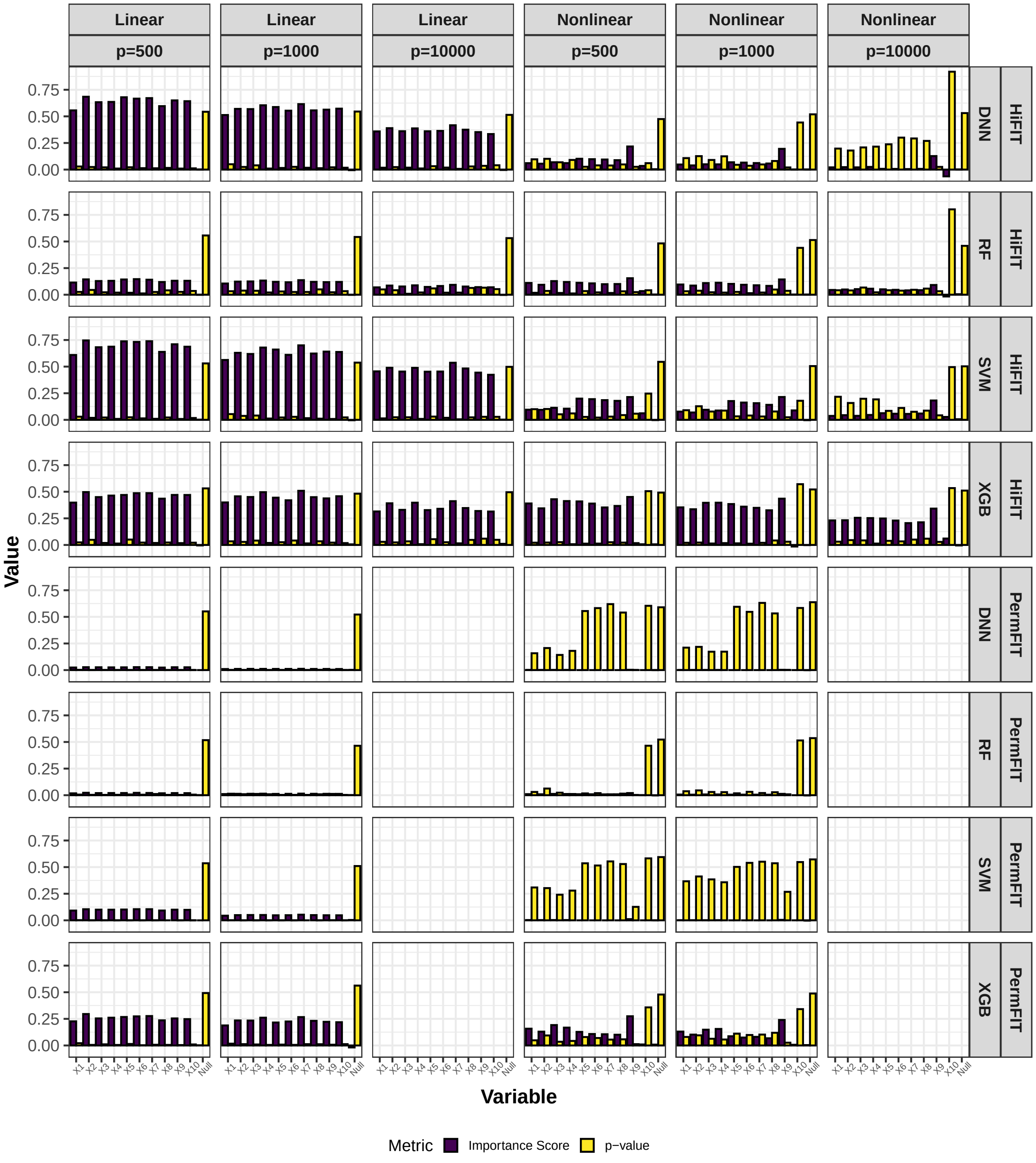


Supplementary Figure 3: **HiFIT Feature Interpretation Results on Binary Outcomes.** Average feature importance scores and p-values for 10 causal variables (denoted as $X_1, ..., X_{10}$) and the feature set of nuisance features (denoted as null) over 100 repetitions. Importance scores of features not selected by HFS are set to zero.

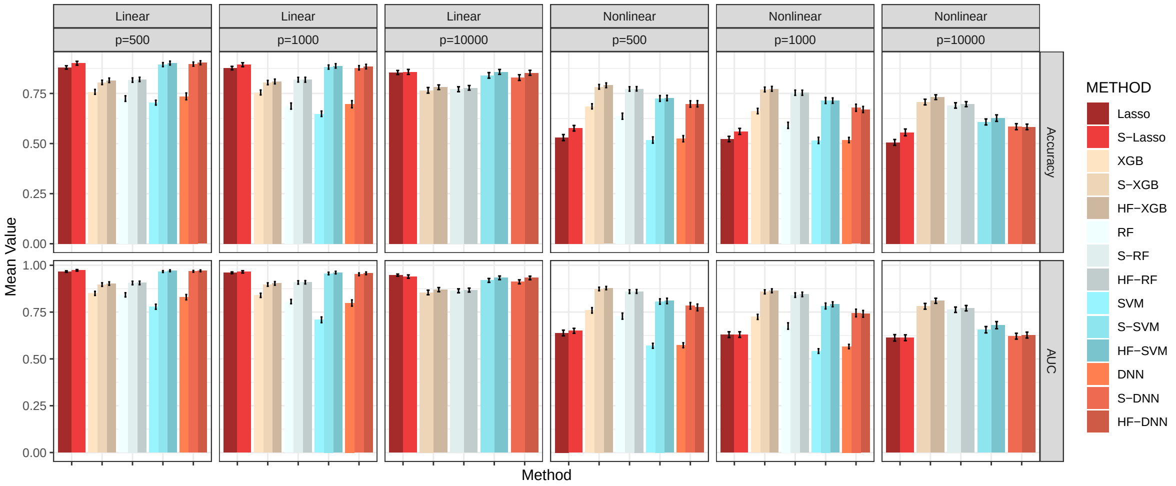


Supplementary Figure 4: **Average Accuracy and AUC for Methods in Comparison on Binary Outcomes.** Lasso, XGB, RF, SVM, and DNN: specific models with all features; S-Lasso, S-XGB, S-RF, S-SVM, S-DNN: specific models with HFS pre-screening; HF-XGB, HF-RF, HF-SVM, HF-DNN: specific models with HiFIT feature selection. Simulation in each scenario is repeated 100 times.
